# Phthalates are detected in the follicular fluid of adolescents and oocyte donors with associated changes in the cumulus cell transcriptome

**DOI:** 10.1101/2024.04.04.588126

**Authors:** Dilan Gokyer, Mary J. Laws, Anna Kleinhans, Joan K. Riley, Jodi A. Flaws, Elnur Babayev

## Abstract

**Purpose:** To investigate follicular fluid (FF) phthalate levels in adolescents undergoing fertility preservation compared to oocyte donors and explore its association with ovarian reserve and cumulus cell gene expression.

**Methods:** 20 Adolescents (16.7 ± 0.6 years old) and 24 oocyte donors (26.2 ± 0.4 years old) undergoing fertility preservation were included in the study. Patient demographics, ovarian stimulation and oocyte retrieval outcomes were analyzed for each group. FF levels of 9 phthalate metabolites were assessed individually and as molar sums representative of common compounds (all phthalates: ƩPhthalates; DEHP: ƩDEHP), exposure sources (plastics: ƩPlastic; personal care products: ƩPCP), and modes of action (anti-androgenic: ƩAA) and compared between the two groups.

**Results:** Follicular fluid ƩPlastic and ƩPCP levels were significantly higher in adolescents compared to oocyte donors (p<0.05). Follicular fluid ƩDEHP, ƩPlastic, ƩPCP, ƩAA, and ƩPhthalates levels were positively associated with antral follicle count (AFC) (p<0.05) in oocyte donors when adjusted for age, BMI, and race/ethnicity. RNA-seq analysis revealed 248 differentially expressed genes (DEGs) in cumulus cells of adolescents within the top quartile (n=4) of FF ƩPhthalates levels compared to the adolescents within the bottom half (n=9). Genes enriched in pathways involved in cell motility and development were significantly downregulated.

**Conclusion:** Adolescents undergoing fertility preservation cycles demonstrate higher levels of phthalate metabolites in their follicular fluid compared to oocyte donors. Phthalate metabolite levels in FF are associated with higher AFC levels in oocyte donors. Higher phthalate levels in FF are associated with alterations in the cumulus cells transcriptome in adolescents.

**Capsule Summary:** Phthalates are detected in the follicular fluid of adolescents and oocytes donors, and the levels are increased in the follicular fluid of adolescents. Higher total phthalate levels in follicular fluid are associated with altered cumulus cells transcriptome in adolescents.

## Introduction

Endocrine-disrupting chemicals (EDCs) mimic, block, or interfere with the hormonal homeostasis [1–4]. Phthalates are ubiquitous environmental contaminants that act as plasticizers in items such as PVC pipes, lubricants, medical tubing, clothing, toys, and food containers to enhance their flexibility and durability [5]. Additionally, they serve as solvents in personal care products (PCPs) and perfumes to stabilize fragrances [5]. These compounds undergo rapid hydrolysis and are metabolized into active monoesters in the body [6]. They are then detected in bodily fluids and subsequently excreted in urine or feces [7]. Alarmingly, the extensive use of phthalates resulted in nearly universal human exposure and over 99% of US women have detectable levels of phthalates in their urine [8]. Women typically exhibit higher levels of phthalates in their biological fluids compared to men of similar ages which could likely be attributed to the prevalent use of these molecules in cosmetics and PCPs [9,10].

Animal studies have demonstrated the impact of phthalates on reproductive health and their potential toxicity on ovaries [11,12]. Phthalates disrupt hormonally mediated processes primarily through interfering with the regulation of hypothalamic-pituitary-gonadal (HPG) axis, consequently altering the levels of gonadotropin releasing hormone (GnRH), and luteinizing hormone (LH) and follicle stimulating hormone (FSH) [13]. In mice, long-term dietary exposure (6 months) to di(2-ethylhexyl) phthalate (DEHP) and diisononyl phthalate (DiNP) has been shown to increase the expression of genes (*Nr5a1*, *Cga*, *Lhb*) that regulate gonadotropin hormones in the pituitary which results in elevated FSH levels and decreased levels of estradiol (E_2_) and LH compared to controls [14]. Additionally, long-term exposure to DEHP and DiNP in mice leads to the disruption of ovarian folliculogenesis, manifested by an increase in the number of primordial follicles and a decrease in the number of preantral and antral follicles [14–16]. Prenatal exposure to DEHP has been shown to induce transgenerational effects in mice ovaries through presumed epigenetic modifications. These modifications result in accelerated folliculogenesis, altered reproductive hormone levels (increased E_2_ and FSH and decreased testosterone and inhibin B), and an increase in ovarian cysts in a multigenerational manner [17,18].

In humans, an elevated urinary concentration of MiBP phthalate correlates with increased odds of premature ovarian insufficiency (POI) [19]. Exposure to DEHP is linked to a decrease in antral follicle count (AFC), a widely accepted marker of ovarian reserve, with a more pronounced effect observed in women under 37 years of age compared to those over 37 [20]. Our group recently demonstrated that urinary phthalate levels are associated with increased ovarian volume in midlife women (45-54 years old). This was associated with higher anti-Müllerian hormone (AMH) and E_2_ levels suggesting the potential folliculogenesis promoting effects of phthalates in this age group [21]. Additional epidemiological studies have established a correlation between EDCs and adverse reproductive and developmental outcomes, including ovulatory irregularities, earlier menarche onset, decreased fertility, decreased semen quality, shortened gestation, higher rate of pregnancy loss [22,23], malformations of reproductive tract such as shorter male anogenital distance [24] and premature breast development in young girls [25].

While urinary phthalate measurements have been the primary method of assessing exposure to EDCs, follicular fluid (FF) provides a more direct estimate of ovarian exposure [26]. Recent studies have highlighted the presence of phthalates in FF and established a direct correlation between exposure to these compounds and detrimental effects on ovaries [27–31]. Elevated concentrations of FF phthalate metabolites in women undergoing infertility treatment are associated with reduced pregnancy rates in Denmark [29]. Similarly, a study from China associated elevated FF phthalate levels with decreased intrafollicular reproductive hormone levels (estradiol, testosterone, progesterone) in women undergoing *In Vitro* Fertilization (IVF), suggesting potential disruptions in granulosa and theca cell endocrine function [27].

There is limited knowledge on phthalate exposure and its effects on reproductive parameters in children and young adults. IVF cycles in post-pubertal adolescents and oocyte donors give a relatively easy access to the immediate oocyte microenvironment, cumulus cells (CCs) and FF. These cells and biofluid offer valuable insights into the quality of the associated gamete and capture crucial changes occurring with advanced reproductive age [31–33]. Cumulus cells support growth and metabolism of the oocyte, facilitate its maturation, and play a crucial role in ovulation and fertilization [32–36]. FF contains hormonal signals and reflects the metabolism, and synthetic capacity of surrounding granulosa and cumulus cells [37–39]. Therefore, the assessment of FF phthalate metabolite levels provides a direct measurement of ovarian phthalate exposure, and the investigation of cumulus cell molecular signature will provide insights into their biological effects.

In this study, our primary objective was to measure the phthalate metabolite levels in FF of adolescents undergoing fertility preservation and compare it with oocyte donors which represent a population with optimal ovarian reserve and egg quality. We also aimed to examine the associations between exposure to different phthalate metabolites and ovarian reserve parameters (e.g., AMH, AFC) as well as IVF outcomes (e.g., number of retrieved mature oocytes (MIIs) and the subsequently generated blastocysts). We then analyzed the association of FF phthalate levels with cumulus cell gene expression.

## Materials and Methods

### Population

We retrospectively analyzed the cumulus cell and follicular fluid samples collected as part of a project aiming to investigate the cumulus cell gene expression and follicular fluid cytokine levels in adolescents compared to oocyte donors at Northwestern Fertility and Reproductive Medicine Center between November 1, 2020, and May 1, 2023. These samples were collected from adolescents (10-19 years old) and oocyte donors (22-30 years old) undergoing ovarian stimulation and oocyte retrieval. There were no exclusion criteria for these participants. Samples were collected from a single ovarian stimulation cycle for each participant. Age, race/ethnicity (as reported by the participant), past medical and surgical history, medication use, results of ovarian reserve and hormone laboratory testing, ovarian stimulation parameters, the information on the number and maturation stage of oocytes, and the number of blastocysts after IVF was collected. All adolescents (N=20) and oocyte donors (N=24) that had FF available in the repository were included in this study.

### Assessment of Follicular Fluid Phthalate Metabolites

Follicular fluid samples were previously collected from the two dominant follicles (18-20 mm) of each ovary per participant, treated with Halt protease inhibitor cocktail (Cat #87786, Thermo Fisher Scientific) to avoid protein degradation, cells were pelleted via centrifugation at 380 × g for 10 min at 4 °C, and the FF supernatant was stored at -80^°^C. These FF samples were used to quantify phthalate metabolites with liquid chromatography-mass spectrometry (LC-MS) by the Carver Metabolomics Core, University of Illinois Urbana-Champaign Roy. J. Carver Biotechnology Center, using previously published methods with modification [21]. Briefly, aliquots of 25 µL follicular fluid were added to 1.5 mL tubes, followed by 5 µL analytical stable isotope-labeled internal standards and 75 µL methanol. Tubes were then vortexed, centrifuged, and had supernatant transferred to LC glass vials. Using an Agilent 1200 HPLC system (Agilent Technologies, Santa Clara, CA, USA), metabolites were separated using a Phenomenex Gemini C6-Phenyl (3µm, 100 x 2mm) column (Phenomenex, Torrance, CA, USA) via a gradient method consisting of mobile phase A: H2O+0.1% formic acid and mobile phase B: acetonitrile +0.1% formic acid. Phthalates levels were reported individually for 9 metabolites (mono-2-ethylhexyl phthalate (MEHP), mono-(2-ethyl-5-hydroxyhexyl) phthalate (MEHHP), mono-(2-ethyl-5-carboxypentyl) phthalate (MECPP), mono-(2-ethyl-5-oxohexyl) phthalate (MEOHP), mono-(3-carboxypropyl) phthalate (MCPP), monobenzyl phthalate (MBzP), monoethyl phthalate (MEP), monobutyl phthalate (MBP), and monoisobutyl phthalate (MiBP)). Metabolites were analyzed with a Sciex 5500 QTRAP MS (Sciex, Framingham, MA, USA) operated in negative electrospray mode with multiple reaction monitoring (MRM). Metabolites were quantified with Sciex Analyst 1.7 software using calibration curves adjusted for labeled internal standards. Limit of quantification was 0.5 ng/mL for MEOHP, MEHHP, MBP, MiBP and MCPP, 2 ng/mL for MECPP and MBzP, 5 ng/mL for MEHP and 10 ng/mL for MEP.

The individual levels were used to approximate participants’ exposure to phthalate parent compounds. Exposure to DEHP was approximated as the molar converted sum of 4 FF metabolites using the following equation: ƩDEHP = (MEHP/278) + (MEHHP/294) + (MEOHP/292) +(MECPP/308). Additional phthalate sums were created based on primary sources of phthalate exposure. Sum of plasticizer (ƩPlastic) and personal care product (ƩPCP) phthalate metabolites were estimated as follows: ƩPlastic = (MEHHP/294) + (MEHP/278) + (MEOHP/292) + (MECPP/308) + (MCPP/252) + (MBzP/256) and ƩPCP =(MEP/194) +(MBP/222) +(MiBP/222). Based on previous experimental and epidemiological studies suggesting anti-androgenic activity in the body for certain phthalate metabolites [40–42], the sum of these metabolites (ƩAA) was calculated as (MEHP/278) + (MEHHP/294) + (MEOHP/292) + (MECPP/308) + (MBzP/256) +(MBP/222) +(MiBP/222). Lastly, all 9 FF phthalate metabolites were molar converted and summed to approximate total phthalate exposure (ƩPhthalates).

### Analysis of cumulus cell transcriptome in relation to follicular fluid phthalate levels

Sixteen adolescents and fifteen oocyte donors had cumulus cell transcriptome data available and were included in this analysis. CC transcriptome represented RNA-seq data from 3 mature-oocyte-associated pooled cumulus cells per participant. CC transcriptome of adolescent participants with high (top quartile) total phthalate exposure (ƩPhthalates) in FF was compared to adolescents with lower FF phthalate levels (bottom half). Similar analysis was performed for 2 different primary sources of phthalate exposure (ƩPlastic and ƩPCP) and for oocyte donors.

RNA-seq read count normalization and differential expression were calculated using DESeq2 which employs the Wald test [43]. The cutoff for determining significantly differentially expressed genes (DEGs) was a False Discovery Rate (FDR)-adjusted p-value less than 0.05 using the Benjamini-Hochberg method. Enrichment analysis was performed using downregulated and upregulated DEGs via Metascape online platform (https://metascape.org/gp/index.html#/main/step1) to identify differences in gene ontology (GO) biological pathways across two groups. Top significantly enriched GO terms were plotted using RStudio version 4.3.1 (R Foundation for Statistical Computing, Vienna, Austria, https://www.R-project.org/).

### Statistical Analysis

The normal distribution of the data was evaluated with the Shapiro-Wilk and Kolmogorov-Smirnov test. Analysis between two groups of continuous variables were performed with unpaired two-sided Student’s t-test or Mann-Whitney U test depending on distribution. Categorical variables were analyzed with Fisher’s exact test or Chi-square test. Spearman nonparametric correlation analysis was performed to evaluate the association between adolescents’ age and FF phthalate levels. Multiple linear regression analysis was conducted to assess the associations between follicular fluid levels of phthalates (ΣDEHP, ΣPlastic, ΣPCP, ΣAA, ΣPhthalates) and ovarian reserve parameters (AMH and AFC), the number of MII oocytes retrieved, and blastocysts generated (donors only) adjusting for age, BMI, and race/ethnicity in both groups. P values < 0.05 were considered statistically significant. Outliers were removed via ROUT method (Q value set at 1%). Both analyses, with and without outliers, are presented. GraphPad Prism version 9.0.1 (Boston, Massachusetts USA, www.graphpad.com) was used for statistical analysis.

## Results

### Demographic and clinical characteristics of participants

Adolescent participants included in this study were on average 10 years younger than oocyte donors (16.7 ± 0.6, 10-19 years old, N=20 vs. 26.2 ± 0.4, 22-30 years old, N=24, p<0.0001) (**Figure 1a,b**). Twelve adolescents were diagnosed with cancer prior to fertility preservation, and eight adolescents had a non-cancer diagnosis, including gender dysphoria (n=6) and severe anemia requiring stem cell therapy (n=2) (**Figure 1c**). Five of twelve of these participants with cancer received chemo-and/or radiotherapy prior to fertility preservation. BMI (24.6 ± 1.4 vs. 24.2 ± 0.5 kg/m^2^, p=0.78) and race/ethnicity were similar across groups. Since donors represent a select group of women with good ovarian reserve, not unexpectedly, AMH levels and AFC were significantly higher in oocyte donors compared to adolescents (AMH: 6.5 ± 0.73 vs. 3.4 ± 0.52 ng/ml, p<0.001 and AFC: 27.7 ± 2.2 vs. 17.3 ± 1.4, p<0.0001) which translated into lower total gonadotropin dose requirement (4092 ± 248 vs. 5539 ± 450 IU, p=0.005) and higher peak estradiol (E_2_) levels (3996 ± 364 vs. 2389 ± 300 pg/mL, p<0.001) in oocyte donors compared to adolescents (**Figure 1a**). Luteal phase start (34.78% vs. 3.23%, p=0.003) and abdominal ultrasound monitoring (65% vs 0%, P<0.0001) were more common for adolescents. Other parameters of ovarian stimulation including the number of retrieved oocytes (MII: 20.4 ± 2.9 vs. 27.4 ± 2.3, p=0.06; MI: 1.4 ± 0.3 vs. 2.6 ± 0.4, p=0.07 and GV: 4.0 ± 1.1 vs. 2.7 ± 0.6, p=0.79) were similar between groups.

**Figure 1.**
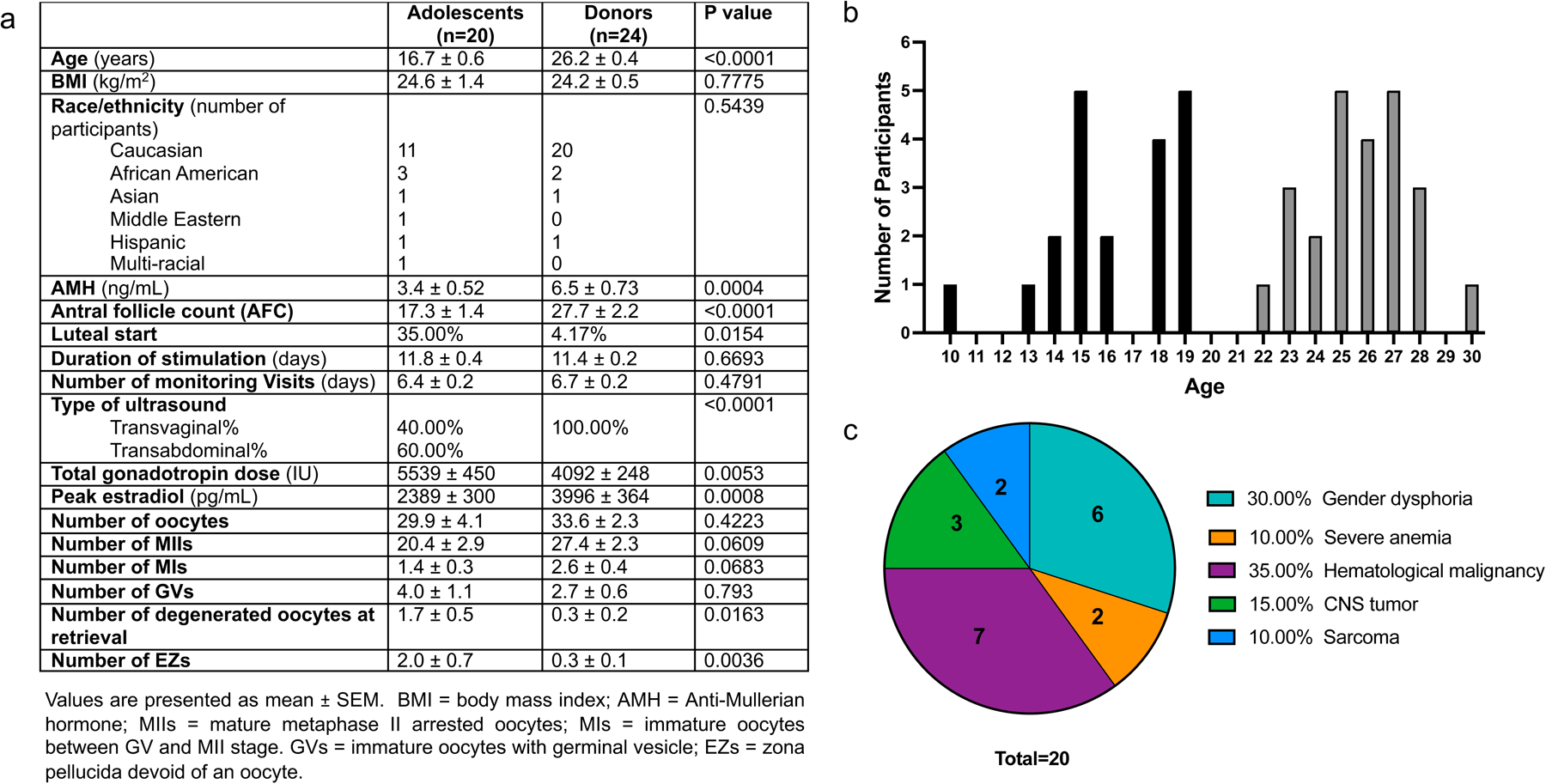
Demographics, IVF cycle characteristics and medical diagnoses of study subjects. (a) Demographics, ovarian reserve, IVF stimulation and outcome parameters, (b) age: adolescents (black bars) and oocyte donors (grey bars) (c) medical diagnoses of adolescents .

### Concentrations of follicular fluid ƩPCP and ƩAA phthalates are higher in adolescents compared to oocyte donors

Out of 9 individual phthalate metabolites, 2 were significantly higher in the FF of adolescents compared to oocyte donors: MBP (27.62 (IQR: 54.00) vs 21.0 (IQR: 249.4) ng/mL, p= 0.01) and MiBP (3.83 (IQR: 11.76) vs. 3.01 (IQR: 105.4) ng/mL, p= 0.04) (**Table 1**). Two phthalate metabolites, MEOHP and MCPP, were absent in the FF of all participants. Personal care product phthalate (ƩPCP: 0.161 (IQR: 0.28) vs 0.123 (IQR: 1.635) pM, p= 0.03), and anti-androgenic phthalate (ƩAA: 0.167 (IQR: 0.287) vs. 0.142 (IQR: 1.673) pM, p=0.045) levels were also significantly higher in FF of adolescents compared to oocyte donors as these two individual metabolites, MiBP and MBP, were included in the calculations of their molar sums (**Table 1**, **Figure 2a,b**). Levels of FF ƩPhthalates, ƩDEHP and ƩPlastic phthalate groups were not significantly different between the two groups (**Figure 2c-e**).

**Figure 2.**
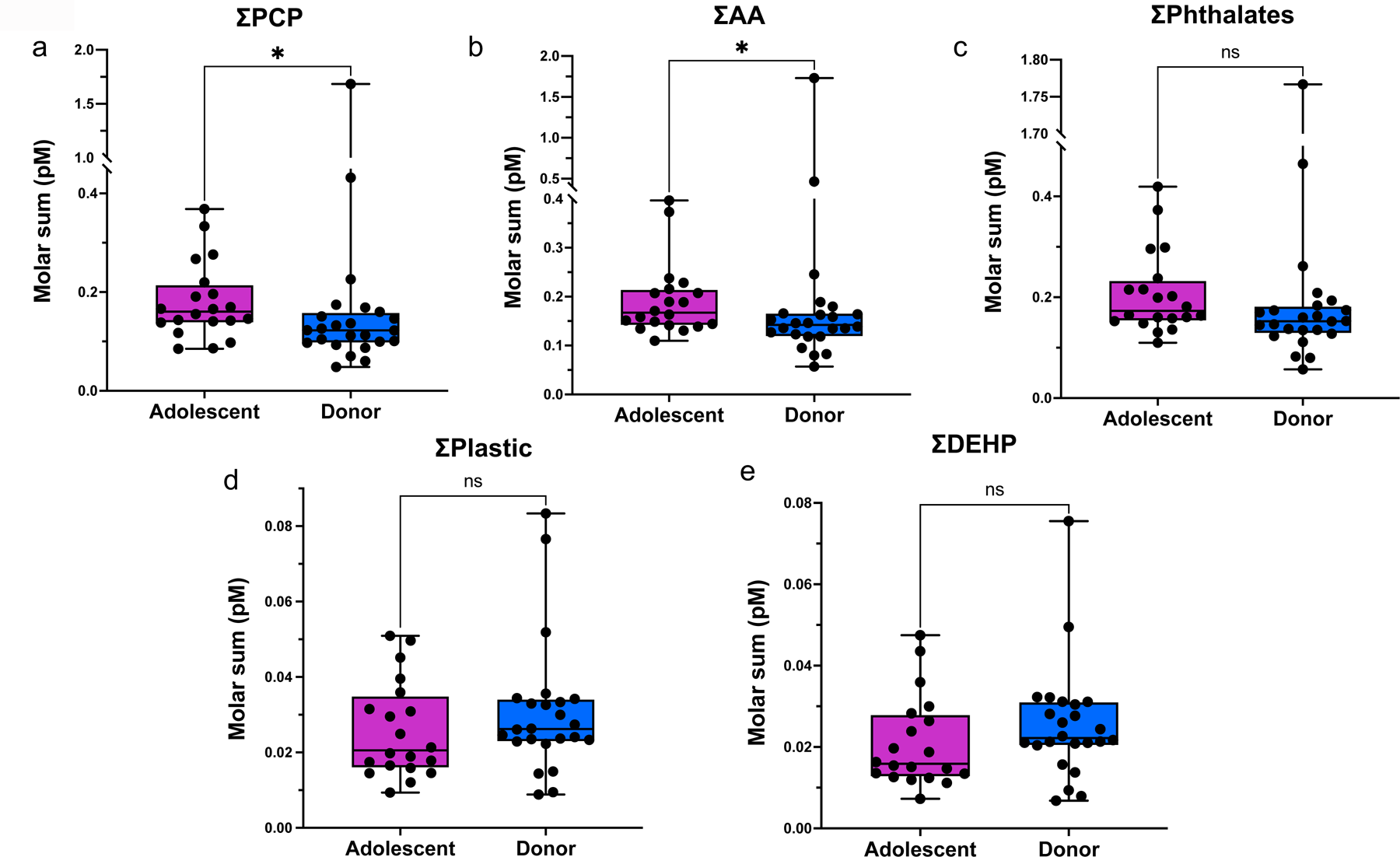
Molar sums of follicular fluid (FF) phthalate metabolites in adolescents compared to oocyte donors. (a) Personal care product (PCP) phthalate metabolites (ƩPCP) and (b) anti-androgenic (AA) phthalate metabolites (ƩAA) were significantly higher in the FF of adolescents compared to oocyte donors. Levels of the sum of (c) all phthalate metabolites (ƩPhthalates), (d) plasticizer phthalate metabolites (ƩPlastic) and (e) DEHP associated phthalate metabolites (ƩDEHP) were not significantly different between the FFs of adolescents compared to oocyte donors. *p ≤ 0.05. DEHP: Di(2-ethylhexyl) phthalate.

**Table 1.**
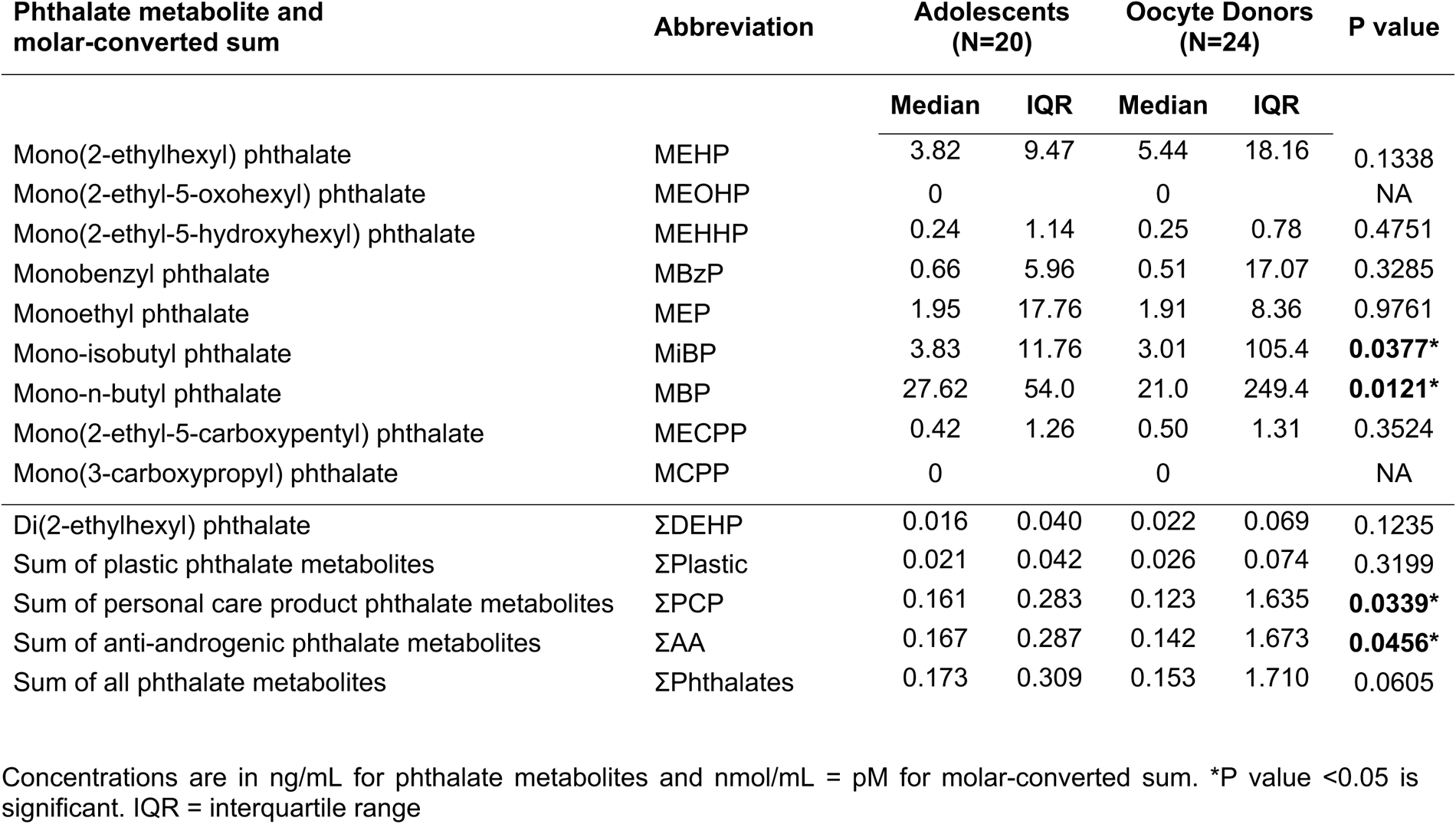
Concentrations of individual phthalate metabolites and their molar sums in follicular fluid of adolescents compared to oocyte donors.

In both groups, some individuals (up to a maximum of 2 per group) exhibited notably different FF phthalate levels compared to others (see individual datapoints in Figure 2). Upon removal of outliers from these datasets, we observed that total phthalates (ƩPhthalates: 0.164 (IQR: 0.189) vs 0.149 (IQR: 0.205) pM, p= 0.03) also demonstrated significantly higher FF levels in adolescents compared to oocyte donors (**Supplementary Table 1, Supplementary Figure 1c**).

Several subgroup analyses were performed to determine whether there is an association between participants’ medical diagnosis, age, racial/ethnic background, and the phthalate levels in their follicular fluid. Our analysis did not reveal any significant differences between FF phthalate levels of adolescents with (n=12) compared to without (n=8) the cancer diagnosis (**Supplementary Figure 2**). Adolescent’s age was also not associated with FF phthalate levels (**Supplementary Figure 3**). FF phthalate levels did not differ in White participants compared to other racial/ethnic backgrounds (Black or Hispanic) in adolescent or oocyte donor groups (**Supplementary Figure 4 and 5**).

### Follicular fluid phthalate levels are associated with antral follicle count in oocyte donors

Multiple linear regression analysis demonstrated that there is no association between FF phthalate levels, and ovarian reserve parameters (AMH, AFC) and the number of mature oocytes retrieved in adolescents when controlled for age, BMI, and race/ethnicity (**Table 2**). However, FF phthalate levels in oocyte donors were positively associated with AFC (p<0.05). There were no associations between FF phthalate levels and AMH, the number of retrieved MII oocytes or generated blastocysts in the oocyte donor group (**Table 3**).

**Table 2.** Association of FF phthalate levels with blood Anti-Mullerian hormone (AMH) levels, Antral follicle count (AFC) and the number of retrieved mature oocyte numbers (MII) in adolescents.

| Independent Variable | Dependent Variable | Parameter Estimate | 95% Confidence Interval |  | p |
| --- | --- | --- | --- | --- | --- |
|  |  |  | LB | UB |  |
| DEHP | AMH | 8.804 | -5.203 | 22.81 | 0.1888 |
|  | AFC | 7.929 | -30.85 | 46.71 | 0.6499 |
|  | MII | 61.42 | -1.281 | 124.1 | 0.0539 |
| Plastic | AMH | 8.603 | -5.108 | 22.31 | 0.1895 |
|  | AFC | 7.187 | -31.31 | 45.69 | 0.6782 |
|  | MII | 57.50 | -6.555 | 121.6 | 0.0729 |
| PCP | AMH | 8.361 | -4.809 | 21.53 | 0.1848 |
|  | AFC | 6.418 | -31.64 | 44.47 | 0.7075 |
|  | MII | 54.48 | -11.24 | 120.2 | 0.0935 |
| AA | AMH | 7.991 | -5.354 | 21.34 | 0.2086 |
|  | AFC | 7.361 | -31.49 | 46.21 | 0.6737 |
|  | MII | 55.53 | -11.43 | 122.5 | 0.0934 |
| Phthalates | AMH | 8.345 | -4.871 | 21.56 | 0.1869 |
|  | AFC | 6.483 | -31.72 | 44.68 | 0.7058 |
|  | MII | 54.81 | -11.35 | 121.0 | 0.0937 |

The analysis was controlled for age, BMI, and race/ethnicity. $p < 0.05$ is significant. LB = lower bound; UB = upper bound; DEHP = Di(2-ethylhexyl)phthalate; PCP = personal care products; AA = anti-androgenic phthalates.

**Table 3.** Association of FF phthalate levels with blood Anti-Mullerian hormone (AMH) levels, Antral follicle count (AFC), the number of retrieved mature oocytes (MII) and generated blastocysts in oocyte donors.

| Independent Variable | Dependent Variable | Parameter Estimate | 95% Confidence Interval |  | p |
| --- | --- | --- | --- | --- | --- |
|  |  |  | LB | UB |  |
| DEHP | AMH | 8.828 | -15.79 | 33.45 | 0.4597 |
|  | AFC | 75.79 | 0.7867 | 150.8 | **0.0479\*** |
|  | MII | 43.69 | -42.64 | 130.0 | 0.2993 |
|  | Blastocyst | 1.131 | -38.56 | 40.82 | 0.9524 |
| Plastic | AMH | 10.04 | -14.04 | 34.11 | 0.3913 |
|  | AFC | 80.50 | 6.822 | 154.2 | **0.0340\*** |
|  | MII | 43.01 | -37.94 | 124.0 | 0.2766 |
|  | Blastocyst | 3.381 | -34.40 | 41.17 | 0.8513 |
| PCP | AMH | 11.40 | -12.83 | 35.63 | 0.3348 |
|  | AFC | 84.88 | 10.49 | 159.3 | **0.0277\*** |
|  | MII | 52.09 | -29.58 | 133.8 | 0.1951 |
|  | Blastocyst | 6.201 | -32.88 | 45.28 | 0.7399 |
| AA | AMH | 11.41 | -12.81 | 35.63 | 0.3341 |
|  | AFC | 84.95 | 10.60 | 159.3 | **0.0275\*** |
|  | MII | 51.88 | -29.71 | 133.5 | 0.1965 |
|  | Blastocyst | 6.254 | -32.70 | 45.21 | 0.7369 |
| Phthalates | AMH | 11.41 | -12.81 | 35.62 | 0.3342 |
|  | AFC | 84.94 | 10.58 | 159.3 | **0.0276\*** |
|  | MII | 51.79 | -29.69 | 133.3 | 0.1966 |
|  | Blastocyst | 6.188 | -32.79 | 45.17 | 0.7398 |

The analysis was controlled for age, BMI, race/ethnicity. p <0.05 is significant. LB = lower bound; UB = upper bound; DEHP = Di(2-ethylhexyl)phthalate; PCP = personal care products; AA = anti-androgenic phthalates.

### Higher follicular fluid phthalate levels are associated with altered cumulus cell gene expression in adolescents

Unsupervised hierarchical clustering and Principal component analysis (PCA) did not demonstrate clear clustering of cumulus cell transcriptomes from adolescents with high (top quartile, n=4) compared to low (bottom half, n=9) FF ƩPhthalates levels (**Supplementary Figure 6**). This is not surprising given the highly differentiated nature of cumulus cells. However, comparative transcriptomic analysis of CCs from adolescents with high compared to low FF ƩPhthalates levels revealed 248 DEGs, with 156 transcripts downregulated and 92 upregulated (**Figure 3a,b**). Genes enriched in biological pathways involved in cell motility and development were significantly downregulated in adolescents with high FF ƩPhthalates levels. Negative regulation of locomotion (GO:0040013, p= 1.1 x 10^-7^), transmembrane receptor protein tyrosine kinase signaling pathway (GO:0007169, p= 4.2 x 10^-6^), growth (GO:0040007, p= 4.6 x 10^-6^), regulation of basement membrane organization (GO:0110011, p= 1.6 x 10^-5^), cell morphogenesis (GO:0000902, p= 1.7 x 10^-5^), positive regulation of cell motility (GO:2000147, p= 2.6 x 10^-5^), actin cytoskeleton organization (GO:0030036, p= 4.6 x 10^-5^), and establishment of cell polarity (GO:0030010, p= 3.8 x 10^-4^) were among the top 20 down-regulated biological pathways (**Figure 3c**). Transcripts in pathways involved in metabolic and catabolic processes were more abundant in CCs of adolescents with high FF ƩPhthalates levels. Xylulose 5-phosphate metabolic process (GO:0051167, p= 5.0 x 10^-7^), small molecule catabolic process (GO:0044282, p= 1.2 x 10^-6^), nucleobase-containing small molecule metabolic process (GO:0055086, p= 1.2 x 10^-5^), glycosyl compound metabolic process (GO:1901657, p= 1.1 x 10^-4^), generation of precursor metabolites and energy (GO:0006091, p= 2.2 x 10^-4^), execution phase of apoptosis (GO:0016050, p= 5.3 x 10^-4^), and reactive oxygen species (ROS) metabolic process (GO:0072593, p= 4.0 x 10^-3^) were among the top 11 up-regulated biological pathways (**Figure 3d**). No DEGs were detected in CCs of oocyte donors with high and low FF ƩPhthalates levels.

**Figure 3.**
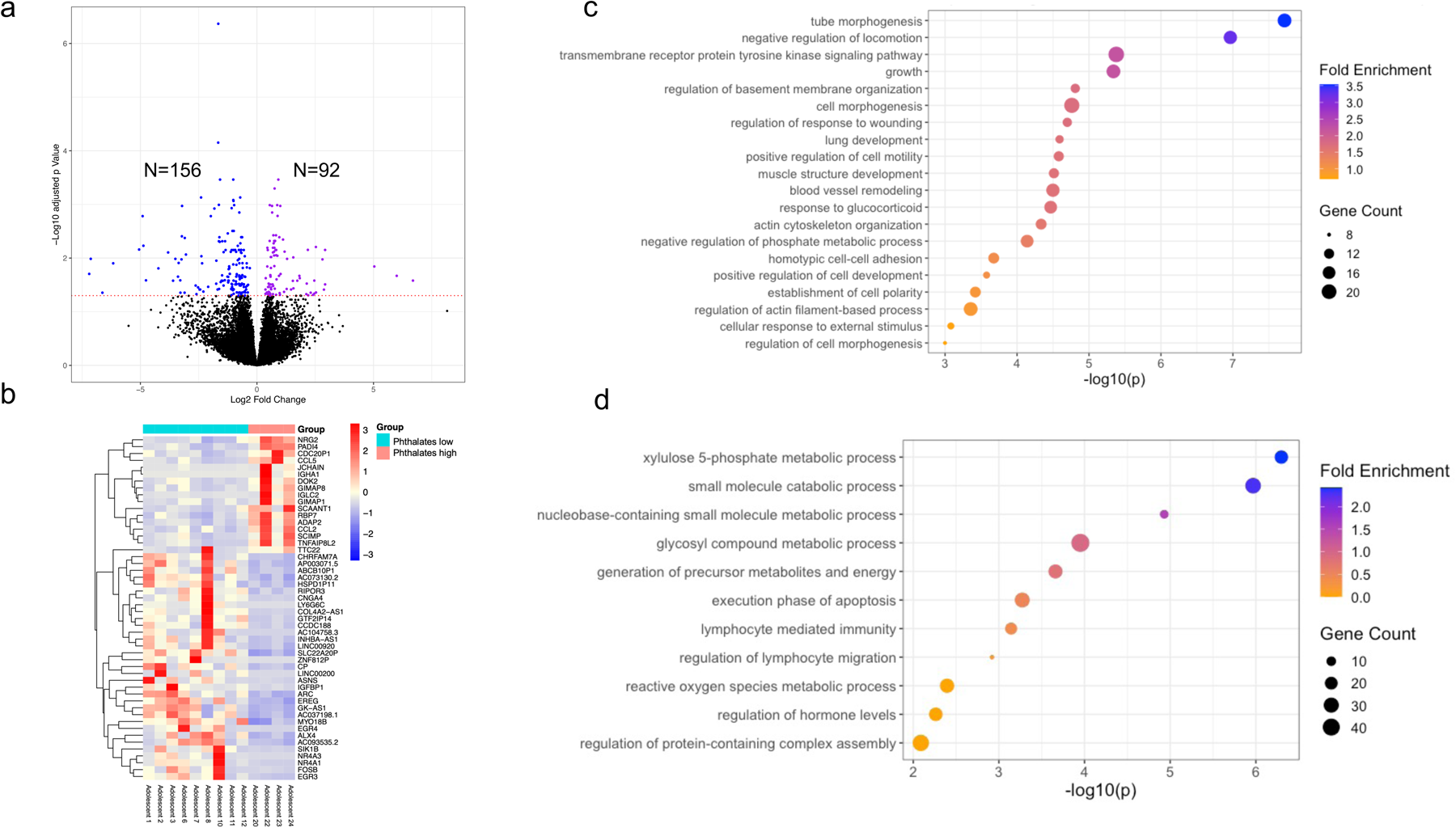
Comparative RNA-seq analysis of cumulus cells (CCs) collected from adolescents with high (top quartile) and low (bottom half) follicular fluid (FF) phthalate levels. a) Volcano plot with downregulated (n=156) and upregulated (n=92) DEGs in adolescents with high compared to low FF ƩPhthalates levels (sum of all phthalates) (dashed red line - adjusted p<0.05) b) Heatmap with the top 50 DEGs with the highest fold change between adolescents with high compared to low FF ƩPhthalates levels c) Top significantly downregulated biological pathway GO terms in adolescents with high compared to low FF ƩPhthalates levels d) Top significantly upregulated biological pathway GO terms in adolescents with high compared to low FF ƩPhthalates levels Dashed red line in volcano plots - adjusted p<0.05. DEGs: differentially expressed genes.

We performed several additional analyses and similarly compared CCs transcriptome in adolescent and oocyte donor groups with high and low ƩPCP and ƩPlastic FF phthalate levels. Two of these revealed DEGs, with fewer genes compared to FF ƩPhthalates analysis. CCs of adolescents with high (top quartile, n=4) compared to low (bottom half, n=9) FF ƩPCP levels revealed 26 downregulated and 16 upregulated DEGs (**Supplementary Figure 7a**). CCs of oocyte donors with high compared to low ƩPCP levels in associated FFs revealed 22 downregulated and 7 upregulated DEGs (**Supplementary Figure 7b**). No significant enrichment in biological pathways was detected for these analyses.

## Discussion

In recent decades, growing concerns have emerged regarding the impact of exposure to environmental chemicals on reproductive health. Research studies in both animals and humans highlighted the adverse effects of phthalates on ovaries [14,21,29]. While studies have shown the associations between higher phthalate levels and poor reproductive parameters such as lower AFC, altered steroid hormone levels (decreased E_2_, increased FSH), and reduced success rates in IVF [19,29], there is limited understanding of the effects of phthalate exposure on reproductive outcomes in adolescents and young adults. In present study, we demonstrated that mean molar sum of FF PCP and AA phthalates (ƩPCP and ƩAA) were significantly higher in adolescents compared to oocyte donors. Multiple linear regression analysis revealed that follicular fluid levels of ΣDEHP, ΣPlastic, ΣPCP, ΣAA, and ΣPhthalates were positively associated with antral follicle count (AFC) in oocyte donors after adjusting for age, BMI, and race/ethnicity. Additionally, RNA-seq analysis identified 248 differentially expressed genes (DEGs) in cumulus cells of adolescents with follicular fluid ΣPhthalates levels within the top quartile compared to those in the bottom half.

Oocyte donors are typically healthy individuals, 21-30 years old, which represent the group with optimal egg quality. The primary distinction between oocyte donors and adolescents in our study was the mean age difference of 10 years. Our analyses revealed FF levels of 2 out of 9 phthalate metabolites, MBP and MiBP, were higher in adolescents. These 2 individual metabolites were included in the calculations of molar sums of ƩPCP, ƩPhthalates and ƩAA, which were also significantly higher in adolescents (for ƩPhthalates only when outliers were removed). The observed phenotype could be attributed to several factors, including the higher frequency of personal care products and perfume usage, differences in clothing choices, dietary habits, and potentially underlying factors such as a cancer or gender dysphoria diagnosis in adolescents as well as different rates of metabolism. Future studies are needed to investigate the association of FF phthalate levels with lifestyle choices in the younger age. The age-dependent rate of phthalate metabolism is also an interesting area for future investigations.

We performed several subanalyses to understand if there is a particular group of individuals among adolescents or oocyte donors that drives this difference. We speculated that adolescents with a cancer diagnosis might have been exposed to phthalates more through medications or medical tubing [44]. Conversely, adolescents with gender dysphoria or who are relatively older could have been exposed to phthalates more through increased use of personal care products (PCPs) or perfumes compared to others [45,46]. However, we found no significant differences in follicular fluid phthalate levels between adolescents with and without a cancer diagnosis. Furthermore, the age of adolescents was not associated with follicular fluid phthalate levels. Previous studies have shown that Black women tend to have significantly higher phthalate exposure than

White women, potentially due to the phthalate content in personal care products (PCPs) preferred by Black women [47,48]. In our study, however, we did not observe any difference in follicular fluid phthalate levels between White participants and those of other races (Black or Hispanic). It is important to note that this lack of difference may be attributed to the low number of individuals in the non-White racial groups (Black or Hispanic) (n=4 in adolescents and n=3 in oocyte donors).

To identify the effects of phthalate exposure on the reproductive outcomes among reproductively young individuals, we examined the associations between FF phthalate levels and ovarian reserve (e.g., AMH, AFC) as well as IVF outcomes (e.g., number of retrieved mature oocytes (MIIs) and the subsequently generated blastocysts) in each group. FF levels of ΣDEHP, ΣPlastic, ΣPCP, ΣAA, and ΣPhthalates were positively associated with antral follicle count (AFC) in oocyte donors after adjusting for age, BMI, and race/ethnicity. These results correlate with our group’s recent report that higher urinary phthalate levels are associated with higher ovarian volume and higher blood AMH/estradiol levels which suggest potential folliculogenesis-promoting properties of these EDCs [15,21].

One limitation of our study is the relatively small sample size. Another limitation is that concomitant exposure to other environmental chemicals cannot be completely excluded, and we were unable to control the analysis for diet and other lifestyle choices [28,49]. In addition, it is important to note that AFC measurements in IVF clinics can be subjective, and in the case of oocyte donors, there may have been an overabundance of follicles, leading to potential inaccuracies in counting by sonographers. Furthermore, the higher utilization of transabdominal ultrasound for AFC measurement in adolescents may have limited accuracy compared to transvaginal ultrasound used in all oocyte donors, as the latter provides clearer visualization of the ovaries with greater detail [50]. These factors could introduce variability and potential bias in AFC measurements.

The immediate microenvironment, CCs and FF, surround the oocyte, and reflect the quality of the associated gamete [51]. The successful release of the oocyte from the ovary during ovulation relies on the transformation of cumulus cells within the expanded cumulus-oocyte complex (COC) into an adhesive, motile, and invasive phenotype [52]. Down-regulated expression of pathways associated with cell migration in adolescent cumulus cells (CC) with high FF ƩPhthalate levels may indicate disordered CC migration due to phthalate exposure, and the upregulation of metabolic and catabolic pathways may signify a compensatory mechanism to eliminate the effects of high FF phthalate metabolites on the CC biology. The mechanisms of phthalate-induced gene expression changes in human cumulus cells is an interesting area of future research. Further studies should investigate why some phthalate parent compounds in FF were associated with the alterations in CC transcriptome while others were not. These studies will shed light on the mechanisms of phthalates’ impacts on human follicle biology and may inform mitigating strategies.

To our knowledge, this is the first study that investigated the phthalate exposure in very young population, adolescents and oocyte donors, and analyzed its effects on cumulus cell biology. Our data shows that phthalates are detected in the follicular fluid at a younger age, and higher levels are associated with altered cumulus cell transcriptome. Our findings pave the way for future investigations assessing the effects of phthalates on human fertility across reproductive lifespan.

## Acknowledgements

This work was supported by the Northwestern University NUSeq Core Facility. We acknowledge Michael La Frano from The Carver Metabolomics Core of the University of Illinois Urbana-Champaign Roy J. Carver Biotechnology Center for their support.

## Declarations

### Funding

This project was supported by Friends of Prentice organization SP0061324 (E.B.), NIH/NICHD K12 HD050121 (E.B.), and NIH R01 ES0341102 (J.A.F.).

### Conflict of interest

The authors have declared that no conflict of interest exists.

### Ethics approval and consent to participate

All participants gave written informed consent according to the protocol approved by Northwestern University Institutional Review Board (STU00213161).

### Availability of data and materials

High-throughput sequencing data will be deposited in Gene Expression Omnibus (GEO) before publication, and the GEO accession number will be provided here.

### Code availability

Not applicable.

### Authors’ contribution

E.B., and J.A.F. conceived the original idea. E.B. designed the experiments. D.G., M.J.L., A.K., J.K.R., and E.B. carried out the experiments. D.G., J.K.R., J.A.F., and E.B. analyzed and interpreted the data. D.G. and E.B. wrote the manuscript. J.K.R. and J.A.F. provided critical discussion, reviewed, and revised the manuscript.

## Notes

### Competing Interest Statement

The authors have declared no competing interest.

