## Supplementary Figure for "Phthalates are detected in the follicular fluid of adolescents and oocyte donors with associated changes in the cumulus cell transcriptome"

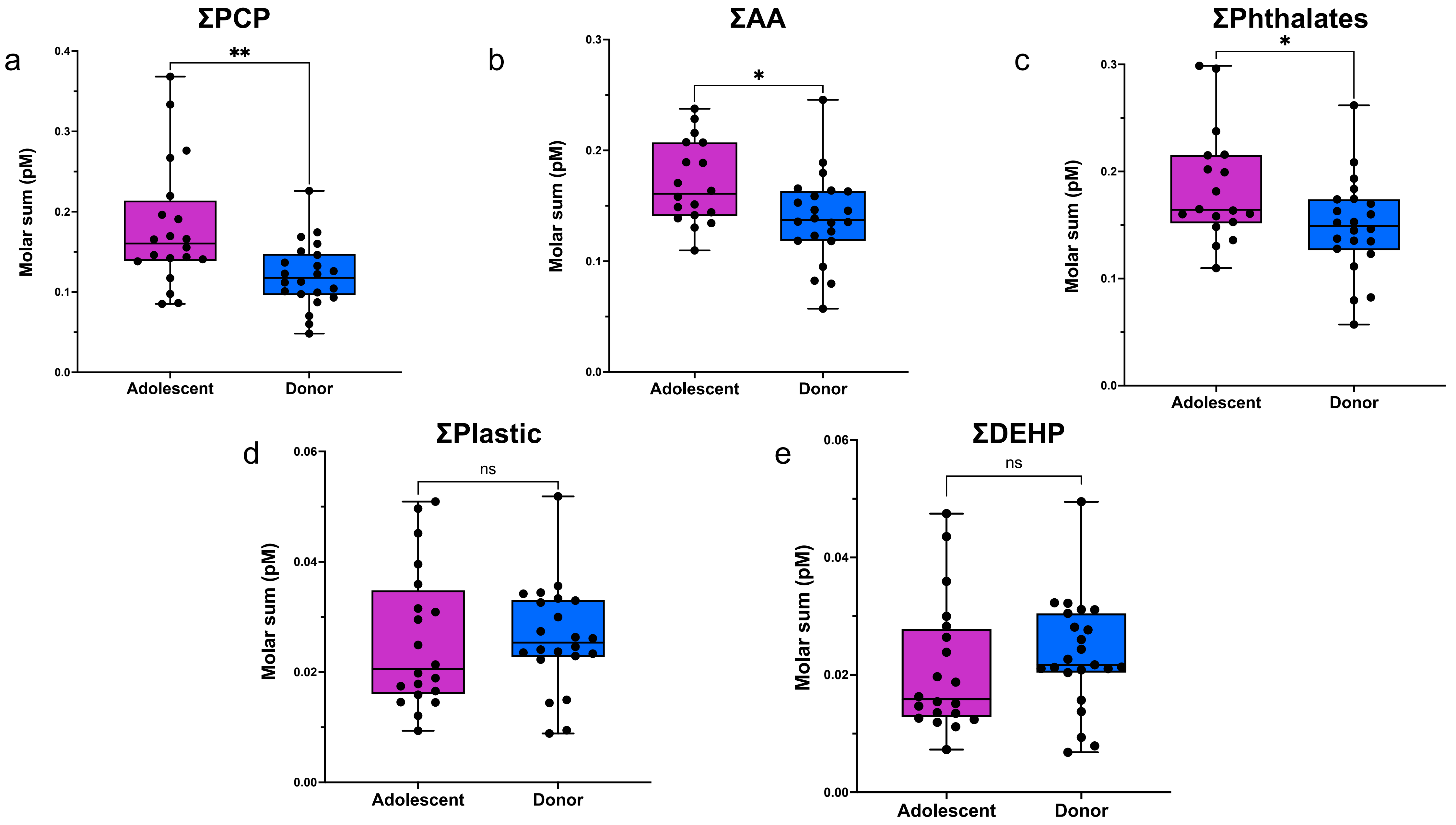


**Supplementary Figure 1. Molar sums of follicular fluid (FF) phthalate metabolites in adolescents compared to oocyte donors.** Each dot represents a patient and outliers were removed from the dataset. *p ≤ 0.05 and ** p ≤ 0.01. PCP: personal care product phthalate, AA: anti-androgenic phthalate, DEHP: Di(2-ethylhexyl) phthalate.

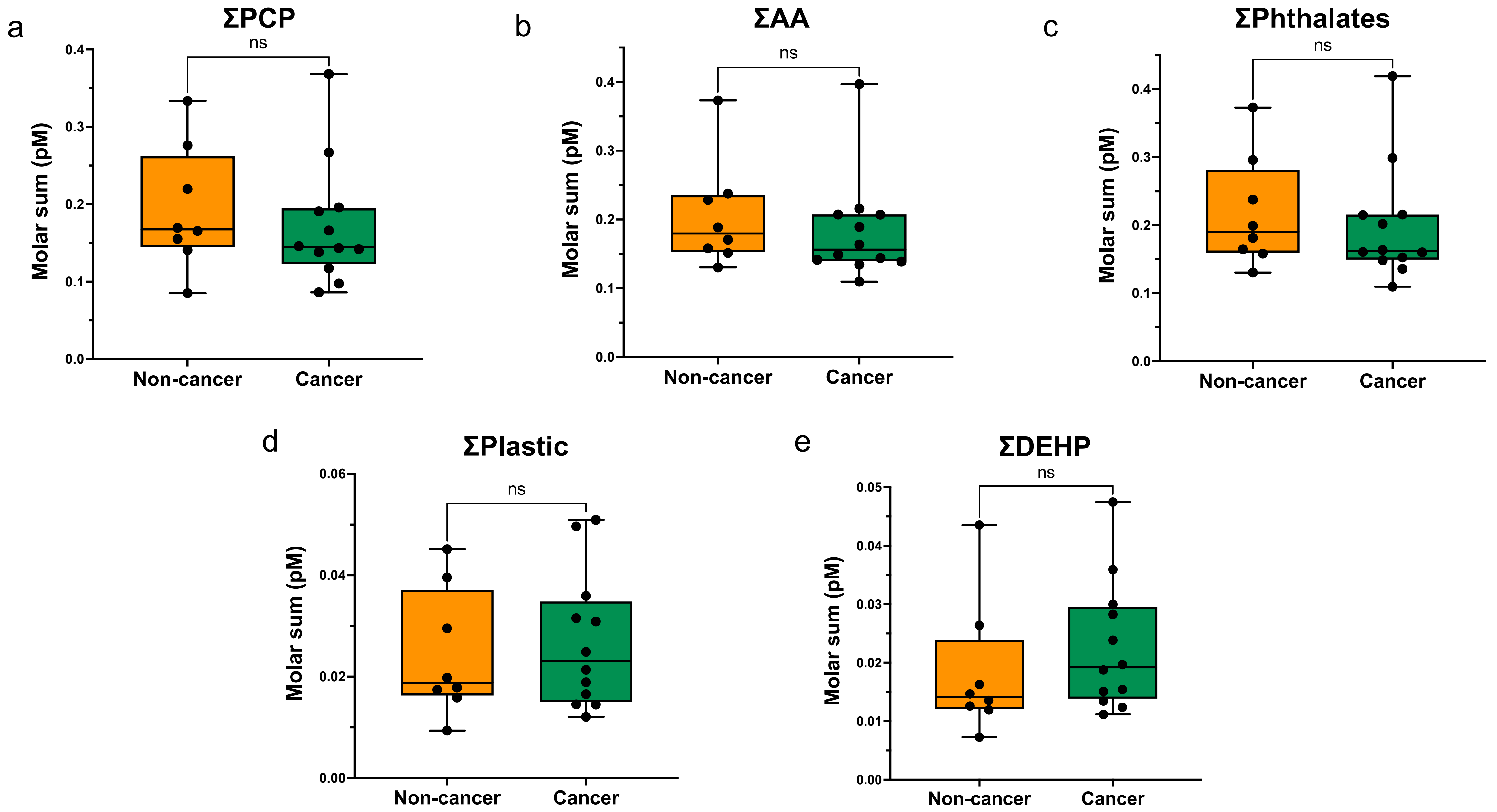


**Supplementary Figure 2. Molar sums of follicular fluid (FF) phthalate metabolites in adolescents with cancer compared to without cancer diagnosis.** Each dot represents a patient. PCP: personal care product phthalate, AA: anti-androgenic phthalate, DEHP: Di(2-ethylhexyl) phthalate.

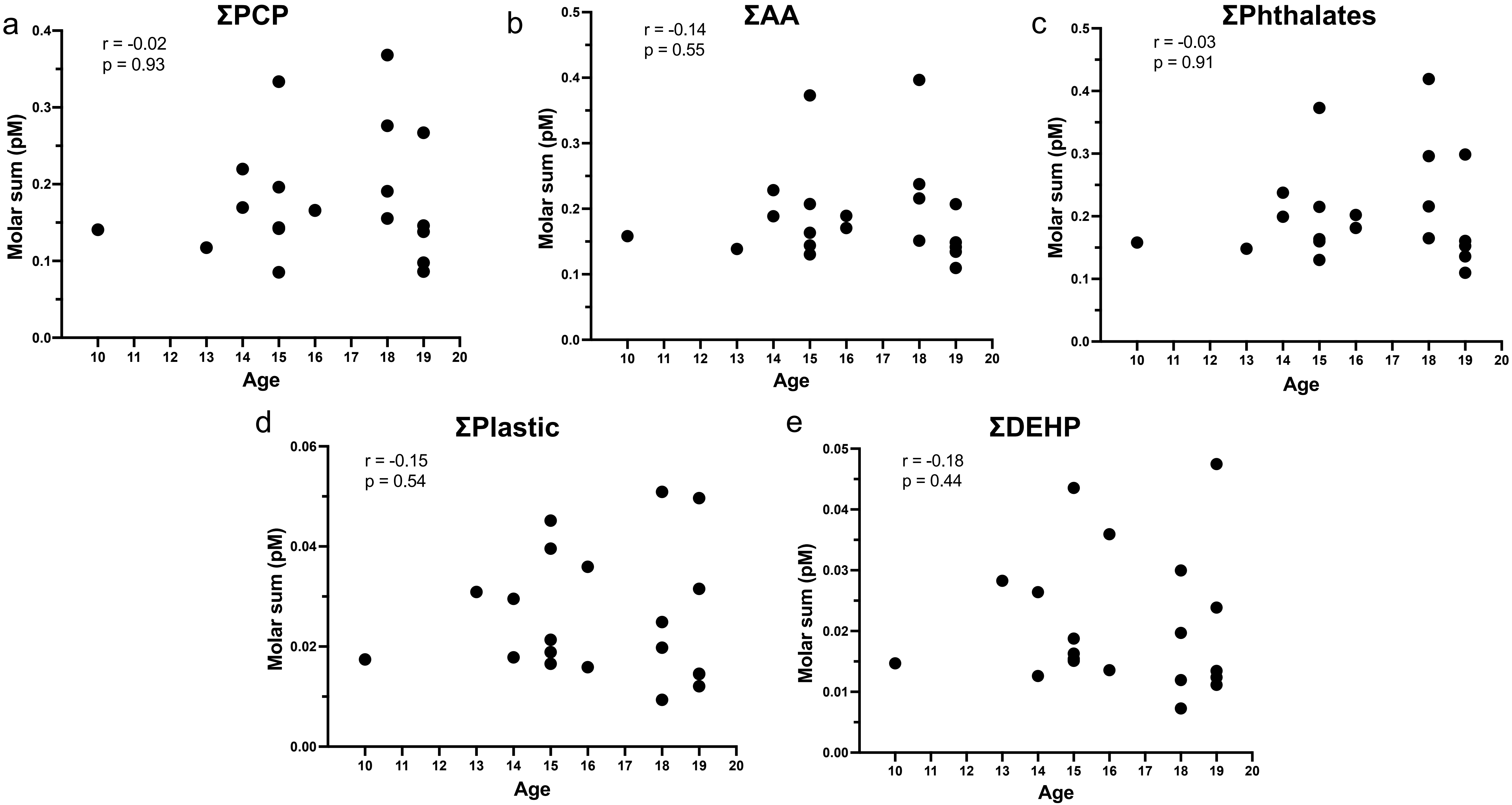


**Supplementary Figure 3. Molar sums of follicular fluid (FF) phthalate metabolites in relation to adolescent's age.** Each dot represents a patient. PCP: personal care product phthalate, AA: anti-androgenic phthalate, DEHP: Di(2-ethylhexyl) phthalate.

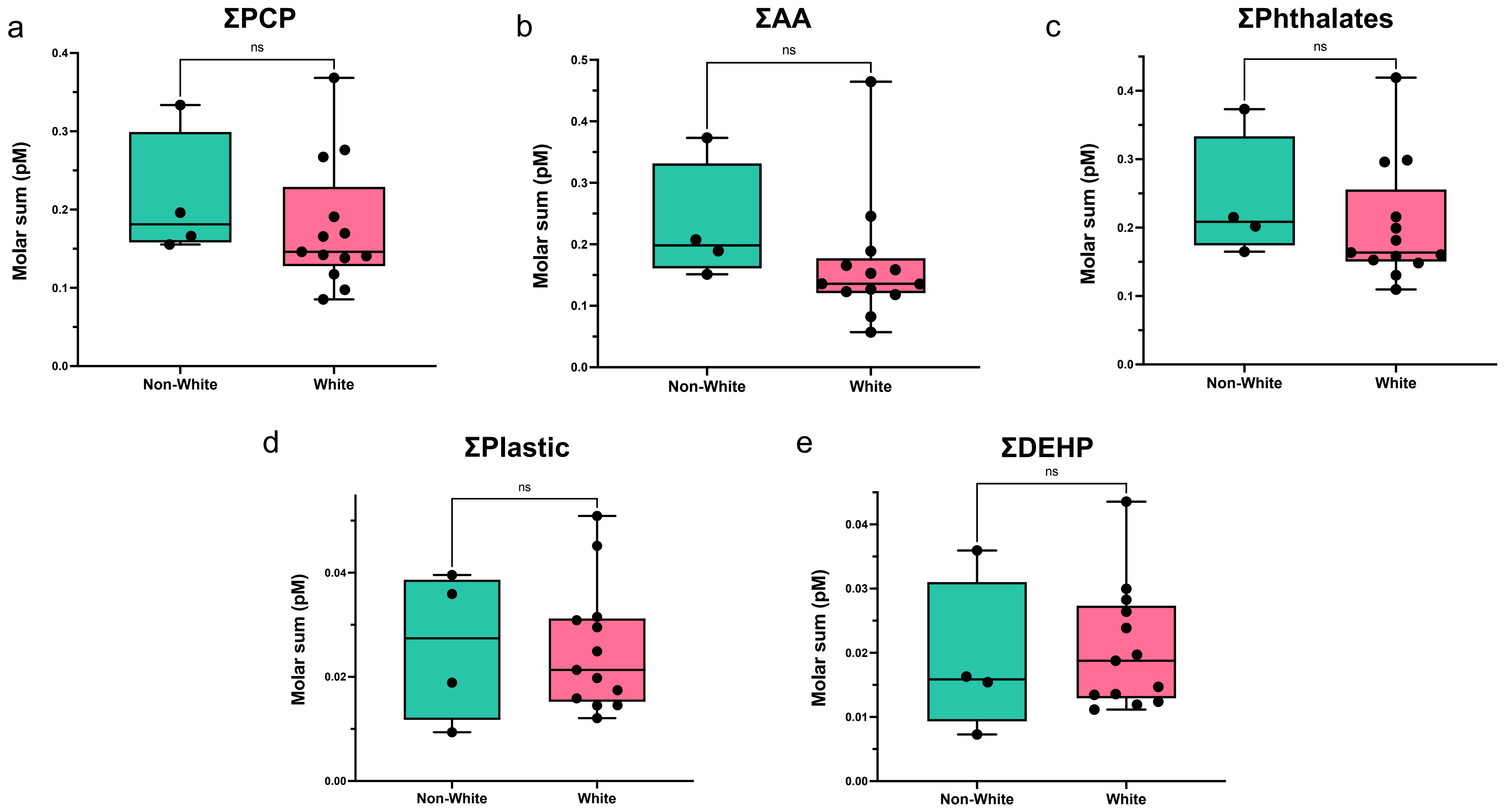


**Supplementary Figure 4. Molar sums of follicular fluid (FF) phthalate metabolites in adolescents with White compared to other racial/ethnic backgrounds.** Each dot represents a patient. PCP: personal care product phthalate, AA: anti-androgenic phthalate, DEHP: Di(2-ethylhexyl) phthalate.

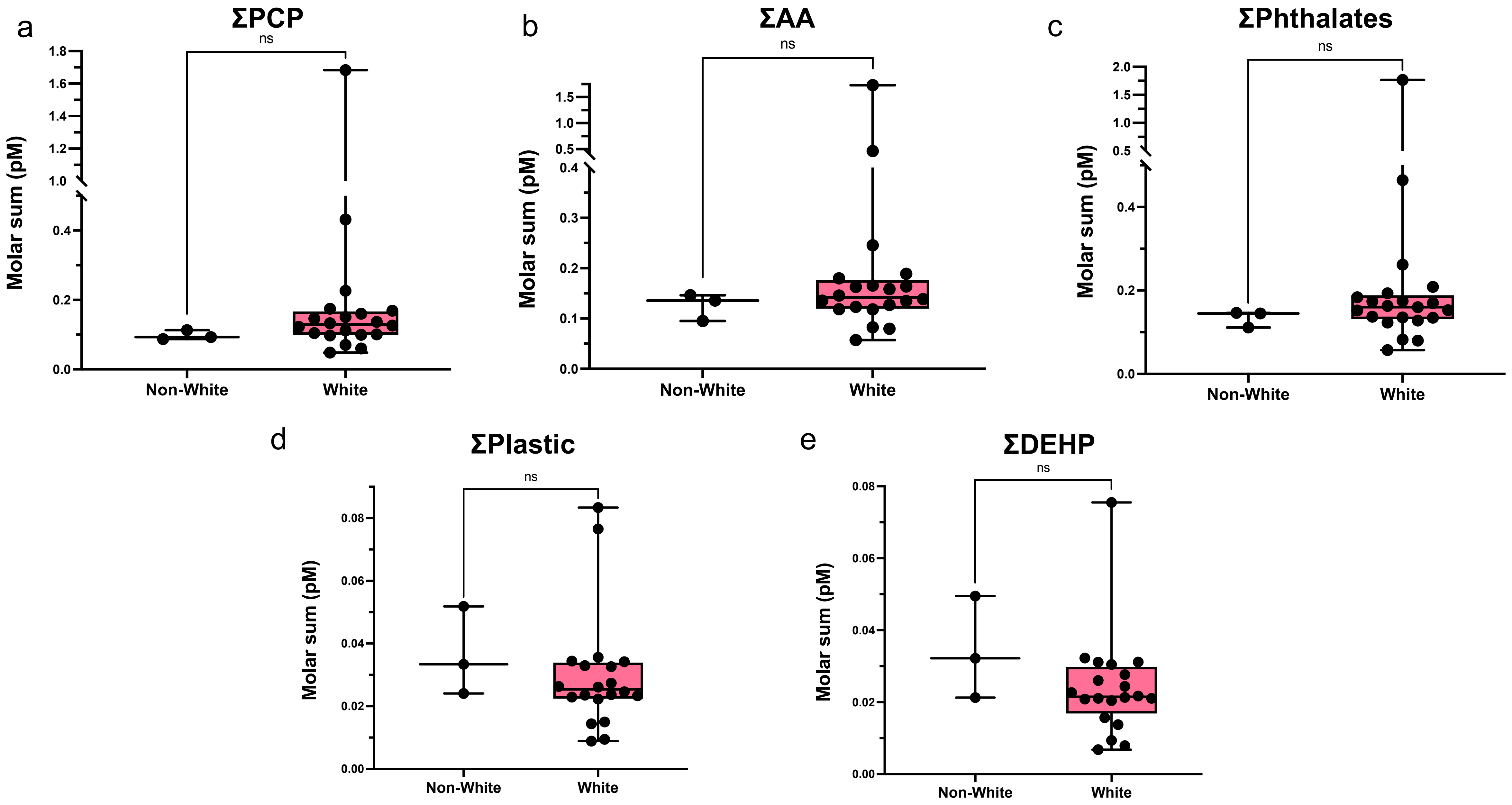


**Supplementary Figure 5. Molar sums of follicular fluid (FF) phthalate metabolites in oocyte donors with White compared to other racial/ethnic backgrounds.** Each dot represents a patient. PCP: personal care product phthalate, AA: anti-androgenic phthalate, DEHP: Di(2-ethylhexyl) phthalate.

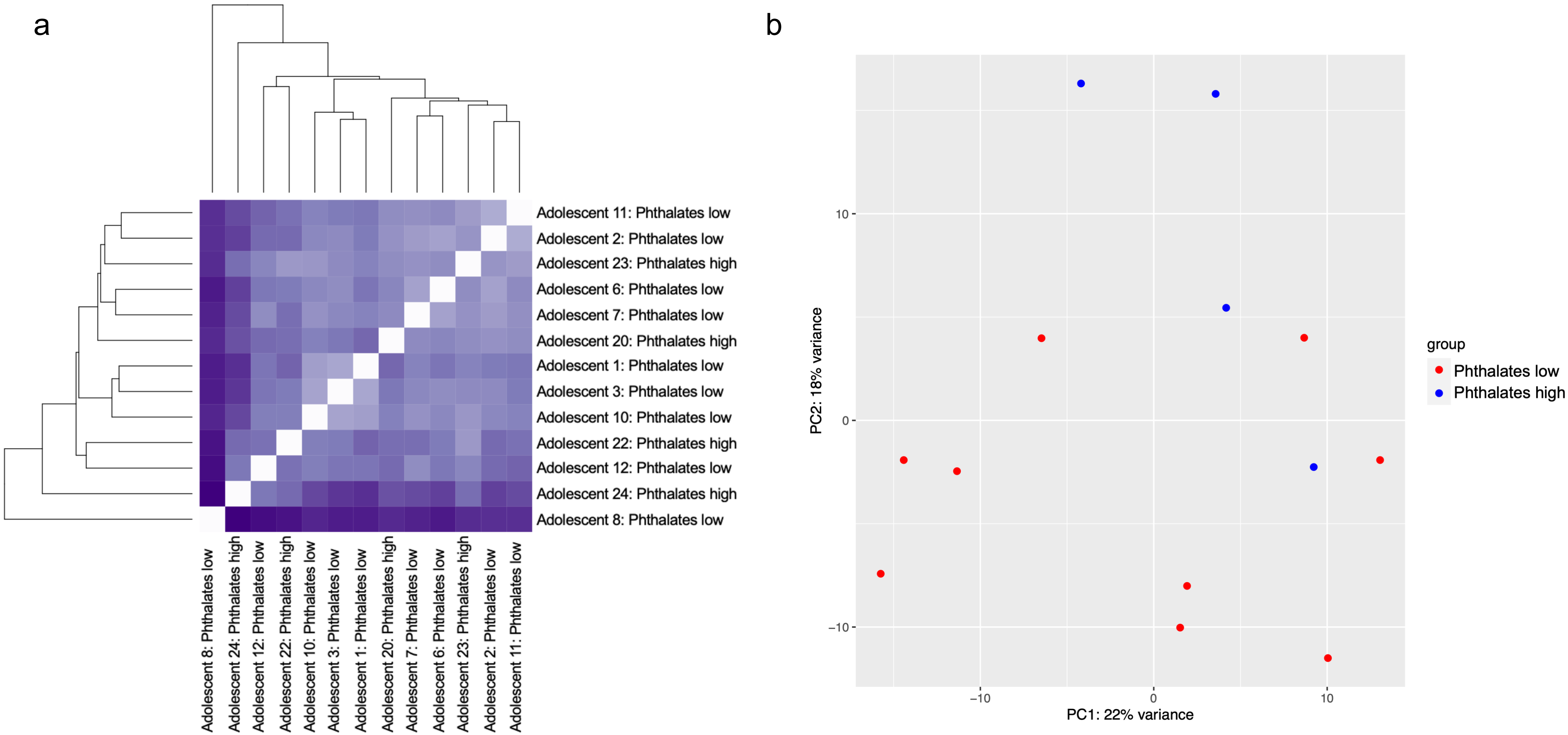


**Supplementary Figure 6. Comparative RNA-seq analysis of cumulus cells collected from adolescents with low (bottom half) and high (top quartile) follicular fluid levels of ΣPhthalates** a) Unsupervised hierarchical clustering b) Principal component analysis. ΣPhthalates: molar sum of all phthalate metabolites

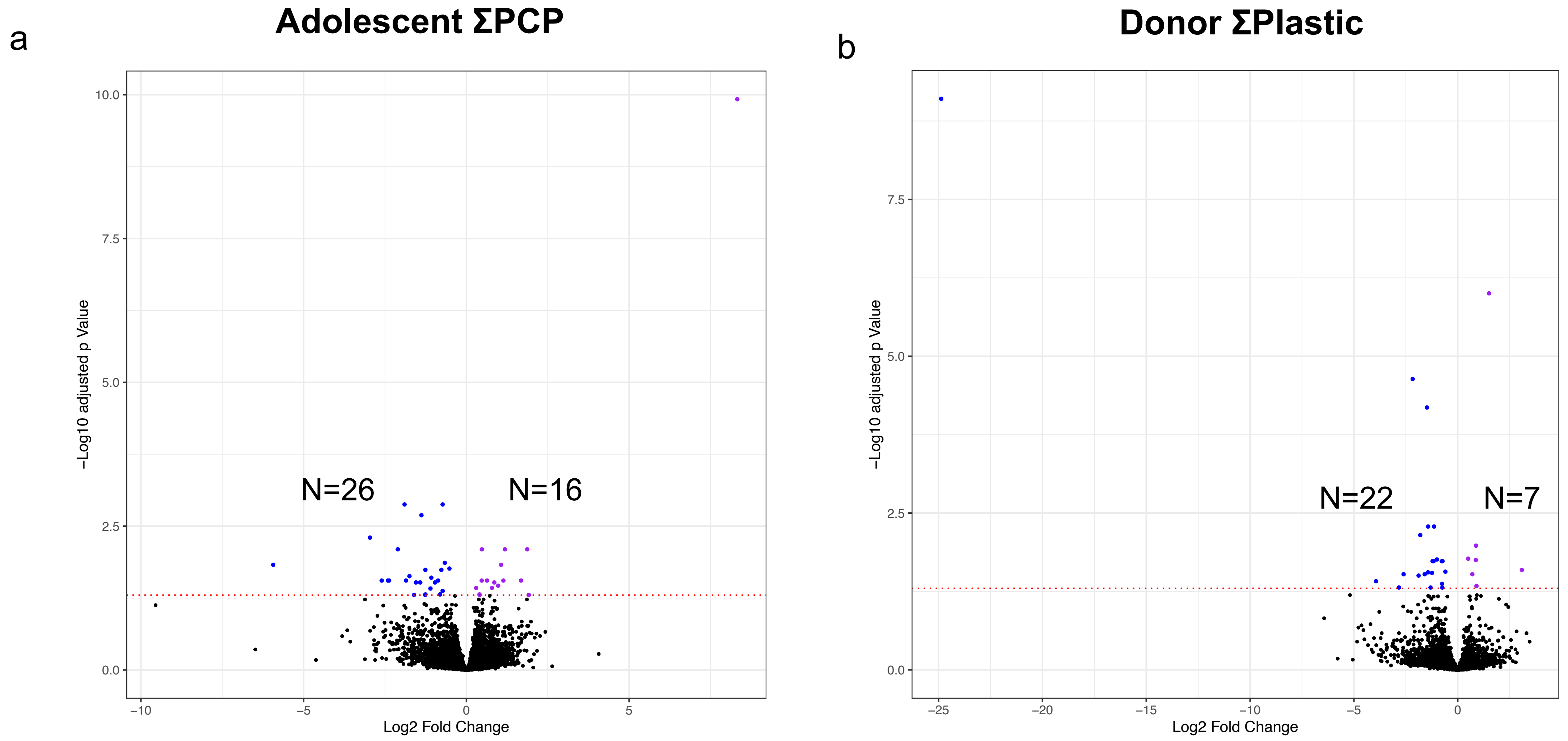


**Supplementary Figure 7. Comparative RNA-seq analysis of cumulus cells (CCs) collected from participants with high (top quartile) and low (bottom half) follicular fluid (FF) phthalate levels** a) Volcano plot with downregulated (n=25) and upregulated (n=16) DEGs in adolescents with high compared to low FF levels of ΣPCP (sum of personal care product) phthalate metabolites b) Volcano plot with downregulated (n=22) and upregulated (n=7) DEGs in oocyte donors with high compared to low FF ΣPlastic levels (sum of plasticizer phthalate metabolites). Dashed red line in volcano plots - adjusted $p<0.05$. DEGs: differentially expressed genes.
