## Supplementary Table 1 for "Phthalates are detected in the follicular fluid of adolescents and oocyte donors with associated changes in the cumulus cell transcriptome"

**Supplementary Table 1. Concentrations of individual phthalate metabolites and their molar sums in follicular fluid of adolescents compared to oocyte donors with outliers removed**

| Phthalate metabolite and molar-converted sum | Abbreviation | Adolescents (N=20) | | Oocyte Donors (N=24) | | P value |
|---|---|---|---|---|---|---|
| | | **Median** | **IQR** | **Median** | **IQR** | |
| Mono(2-ethylhexyl) phthalate | MEHP | 3.82 | 9.47 | 5.36 | 10.04 | 0.1925 |
| Mono(2-ethyl-5-oxohexyl) phthalate | MEOHP | 0 | | 0 | | NA |
| Mono(2-ethyl-5-hydroxyhexyl) phthalate | MEHHP | 0.24 | 1.14 | 0.25 | 0.78 | 0.4751 |
| Monobenzyl phthalate | MBzP | 0.63 | 2.01 | 0.51 | 1.04 | 0.4542 |
| Monoethyl phthalate | MEP | 1.82 | 4.36 | 1.91 | 8.36 | 0.5126 |
| Mono-isobutyl phthalate | MiBP | 3.70 | 6.68 | 2.92 | 4.44 | **0.0103*** |
| Mono-n-butyl phthalate | MBP | 27.62 | 54.0 | 20.92 | 34.64 | **0.0042**** |
| Mono(2-ethyl-5-carboxypentyl) phthalate | MECPP | 0.42 | 0.80 | 0.50 | 0.68 | 0.3006 |
| Mono(3-carboxypropyl) phthalate | MCPP | 0 | | 0 | | NA |
| Di(2-ethylhexyl) phthalate | ΣDEHP | 0.016 | 0.040 | 0.022 | 0.043 | 0.1784 |
| Sum of plastic phthalate metabolites | ΣPlastic | 0.021 | 0.042 | 0.026 | 0.043 | 0.5747 |
| Sum of personal care product phthalate metabolites | ΣPCP | 0.161 | 0.283 | 0.118 | 0.179 | **0.005**** |
| Sum of anti-androgenic phthalate metabolites | ΣAA | 0.161 | 0.128 | 0.142 | 0.189 | **0.0237*** |
| Sum of all phthalate metabolites | ΣPhthalates | 0.164 | 0.189 | 0.149 | 0.205 | **0.0341*** |

Concentrations are in ng/mL for phthalate metabolites and nmol/mL = pM for molar-converted sum. Outliers were removed from the dataset during the analysis. *P value <0.05 and **P value <0.01 is significant. IQR = interquartile range
